# The GARP Domain of the Rod CNG Channel’s β1-subunit Contains Distinct Sites for Outer Segment Targeting and Connecting to the Photoreceptor Disc Rim

**DOI:** 10.1101/2020.10.01.322859

**Authors:** Jillian N. Pearring, Jason R. Willer, Jorge Y. Martínez-Márquez, Eric C. Lieu, Raquel Y. Salinas, Vadim Y. Arshavsky

**Affiliations:** Department of Ophthalmology, University of Michigan, Ann Arbor, MI, 48105, USA; Department of Cell and Developmental Biology, University of Michigan, Ann Arbor, MI, 48105, USA; Albert Eye Research Institute, Duke University, Durham, NC, 27710, USA

**Keywords:** Photoreceptor, cyclic nucleotide-gated channel, traffic, outer segment, cilium

## Abstract

Vision begins when light is captured by the outer segment organelle of photoreceptor cells in the retina. Outer segments are modified cilia filled with hundreds of flattened disc-shaped membranes. Disc membranes are separated from the surrounding plasma membrane and each membrane type has unique protein components. The mechanisms underlying this protein sorting remain entirely unknown. In this study, we investigated the outer segment delivery of the rod cyclic nucleotide-gated (CNG) channel, which is located in the outer segment plasma membrane where it mediates the electrical response to light. We now show that the targeted delivery of the CNG channel to the outer segment requires pre-assembly of its constituent α1 and β1 subunits and that CNGβ1 contains specific targeting information encoded within the glutamic acid-rich region of its N-terminal GARP domain. We also found that the GARP domain connects the CNG channel to photoreceptor disc rims likely through an interaction with peripherin-2 and demonstrated that this function is performed by a proline-enriched region adjacent to the GARP domain. Our data reveal fine functional specializations within the structural domains of the CNG channel and suggest that channel delivery to the outer segment is independent of peripherin-2 interactions.

**Significance Statement:** The precise delivery and organization of signaling proteins in the ciliary outer segment organelle of photoreceptor cells is critical for light detection. We report that the CNG channel, mediating the electrical response to light in rods, contains a region within the N-terminus of its CNGβ1 subunit that encodes the outer segment targeting information for the entire channel. This targeting region is adjacent to a region that connects CNGβ1 to the rims of photoreceptor discs, likely determining the subcellular compartmentalization of the CNG channel into the outer segment plasma membrane.

## Introduction

Photoreceptor cells of the retina are responsible for light detection. They contain a modified primary cilium, called the outer segment, which houses all the molecular components necessary for capturing light and eliciting an electrical response. To increase light sensitivity, outer segments contain a stack of flattened membrane “discs”, so that incoming photons pass through hundreds of membrane surfaces packed with the visual pigment, rhodopsin. Another unique outer segment feature is that it undergoes constant regeneration, which places a heavy demand on the trafficking of signaling and structural proteins to this compartment (Pearring et al., 2013; Goldberg et al., 2016; Spencer et al., 2020b). Defects in these trafficking mechanisms underlie many forms of inherited retinal degeneration, ultimately leading to blindness.

Within the outer segment, there is functional specialization between the membranes of photoreceptor discs and the surrounding plasma membrane. Whereas discs primarily harbor components of the visual signaling pathway (Molday and Molday, 1987; Skiba et al., 2013), the cyclic nucleotide-gated (CNG) channel required to elicit an electrical response to light and the Na/Ca-K exchanger regulating Ca^2+^ dynamics are confined to the plasma membrane (Bauer, 1988; Reid et al., 1990). Previous studies, focused primarily on disc-resident transmembrane proteins, showed that their outer segment trafficking relies on specific sequences for targeting. These proteins include rhodopsin (Sung et al., 1994), peripherin-2 (Tam et al., 2004; Salinas et al., 2013), RDH8 (Luo et al., 2004) and R9AP (Pearring et al., 2014). Two other proteins, GC-1 and PRCD, have been shown to be delivered to the outer segment in a complex with rhodopsin (Pearring et al., 2015; Spencer et al., 2016). Less is known about the outer segment delivery of plasma membrane-resident proteins and even less about their segregation specifically into this membrane.

In this study, we investigated the outer segment targeting and plasma membrane sorting of the CNG channel. This channel is composed of four subunits: three α1 and one β1 (Weitz et al., 2002; Zheng et al., 2002; Zhong et al., 2002). Early studies of CNG channel trafficking, performed by heterologously expressing its subunits in *Xenopus* oocytes, showed that both CNGα1 and the CNGα1/ CNGβ1 complex, but not CNGβ1 alone, are effectively delivered to the oocyte plasma membrane and form functional channels. Yet, only the channel formed by both subunits recapitulates electrical and gating properties of the native channel (Kaupp et al., 1989; Chen et al., 1993; Colville and Molday, 1996; Trudeau and Zagotta, 2002; Zheng et al., 2002). Transgenic expression of the channel in *Xenopus* rods showed that removing the N-terminal glutamic acid-rich protein (GARP) domain from CNGβ1 results in accumulation of the mutant subunit in the biosynthetic membranes and mistrafficking to the plasma membrane of the inner segment (Nemet et al., 2014), suggesting an important role for the GARP domain in CNGβ1 transport.

The CNG channel was reported to associate with the Na/Ca-K exchanger within the outer segment plasma membrane (Molday and Molday, 1998). In addition, the channel is connected to the photoreceptor disc rim through binding to a rim-specific protein, peripherin-2 (Poetsch et al., 2001; Conley et al., 2010). The latter interaction is believed to link the discs with the plasma membrane, likely stabilizing the disc stack within the outer segment.

We now demonstrate that the CNG channel travels through both the ER and Golgi on its route to the rod outer segment and that its specific outer segment delivery requires the channel’s pre-assembly. This targeting is mediated by the glutamic acid-rich region of CNGβ1’s GARP domain. Whereas this targeting region confers outer segment localization, it does not define specific plasma membrane localization. We observed that the adjacent R1-4 region of the GARP domain accumulates in disc incisures likely through peripherin-2 binding. Together, our results show that outer segment trafficking and peripherin-2 interaction are performed by two distinct sites within the CNGβ1 subunit’s GARP domain and suggest that the plasma membrane sequestration of the channel relies on peripherin-2 interaction.

## Materials and Methods

### Animals

Mice were handled following the protocols approved by the Institutional Animal Care and Use Committees at the University of Michigan (registry number A3114-01) and Duke University (registry number A011-14-01). Albino WT CD1 mice used in electroporation experiments were obtained from Charles River. *Cngb1-X1*^*−/−*^ mice (Zhang et al 2011) used in Western blotting and electroporation experiments were kindly provided by Steven Pittler (University of Alabama). Pigmented wild-type C57BL/6J mice used for Western blotting were from Jackson Labs (Bar Harbor, ME). All mice were housed under a 12/12-hour light cycle. The experimenters were not blinded to genotype.

#### In vivo *electroporation*

Retinal transfection of neonatal mice by the *in vivo* electroporation technique (Matsuda and Cepko, 2007) was used with modifications described in (Pearring et al., 2015; Salinas et al., 2017) to express exogenous constructs in mouse rods. DNA constructs were electroporated into the retinas of wild-type CD1 or *Cngb1-X1*^*−/−*^ neonatal mice. Following anesthetization of neonatal mice (P0-P2) on ice, the eyelid and sclera were punctured at the periphery of the eye using a 30-gauge needle. A blunt-end 32-gauge needle was advanced through the puncture wound until reaching the subretinal space, and 0.3–0.5 µl of concentrated plasmid DNA (2 µg/µl of the construct of interest and 1 µg/µl soluble mCherry to visualize transfected cells) was deposited. A tweezer-type electrode (BTX) was placed over the mouse’s head with the positive electrode overlying the injected eye. Five 100V pulses of 50 ms duration were applied using an ECM830 square pulse generator (BTX). Neonates were returned to their mother and allowed to develop until postnatal day 21 when mice were sacrificed by CO_2_ inhalation followed by decapitation and retinal tissue was collected for analysis.

#### Generation of transgenic tadpoles

Transgenic *Xenopus* tadpoles were produced using restriction enzyme-mediated integration developed previously (Kroll and Amaya, 1996; Amaya and Kroll, 1999), with modifications described in (Batni et al., 2000; Whitaker and Knox, 2004).

### Primary antibodies

The following antibodies were generously provided by: Robert Molday, University of British Columbia (mAb 1D1, anti-CNGa1; and mAb 8G8, anti-GARP); Steven Pittler, University of Alabama at Birmingham (pAb, anti-CNGβ1); Gabriel Travis, University of California Los Angeles (pAb, anti-peripherin-2). The polyclonal antibody against Rom-1 was generated in the Arshavsky laboratory (Gospe et al., 2011). Rabbit antibody against R9AP sequence 144–223 is described in (Keresztes et al., 2003). Commercial antibodies were: mAb 1D4, anti-rhodopsin (Abcam, ab5417); pAb M-18, anti-ABCA4 (Santa Cruz, sc-21460); mAb anti-Na/K-ATPase (Santa Cruz, sc-58628); mAb M2, anti-FLAG (Sigma-Aldrich, F1804); pAb, anti-FLAG (Sigma-Aldrich, F7425); pAb, anti-GFP conjugated to Alexa Fluor 488 (Molecular Probes); and pAb, anti-MYC (Cell Signaling, 2278S).

### DNA Constructs

DNA constructs were generated using standard PCR-based subcloning methods. Point mutations were generated using the QuikChange II XL kit (Stratagene). All DNA constructs were cloned between a 5’ AgeI and 3’ NotI sites and sequence confirmed. For mouse *in vivo* electroporation, the pRho plasmid driving gene expression under the 2.2 kb bovine rhodopsin promoter was used (Addgene, plasmid # 11156). For Xenopus transgenics, the XOP1.5 vector was used for constructs containing GFP, while untagged and Myc-tagged CNGβ1 constructs were expressed using a dual promoter strategy described in Baker et al. (2008). In brief, along with the *Xenopus* opsin promoter driving expression of the various CNGβ1 constructs in rods, a cassette containing the γ-crystallin promoter was used to drive GFP expression in the lens. DNA templates were obtained as follows: human *Cngb1* was a gift from Van Bennett (Duke University), mouse *Cnga1* and *Cngb1* were amplified from mouse retina cDNA (Stratagene, La Jolla, CA); the Myc epitope was added using overlap extension PCR. All chimeric constructs were produced using overlap extension PCR. All forward and reverse primers were designed to introduce 5′-AgeI and 3′-NotI sites respectively. A complete list of primers can be found in Table 1.

**Table 1.**
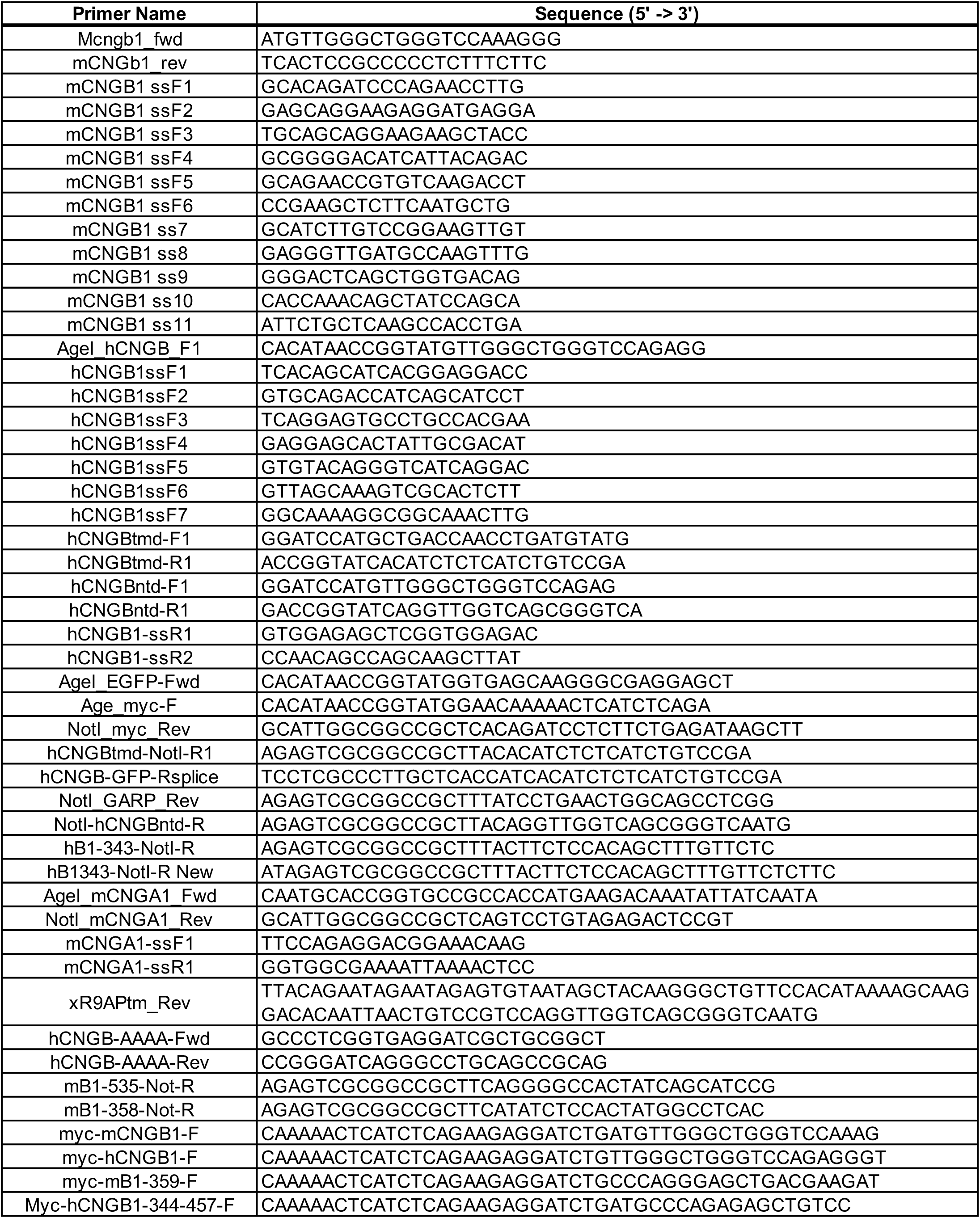

### Immunofluorescence

#### Xenopus retinal cross-sections

GFP expressing transgenic tadpoles at developmental stages 43-45 were anesthetized in 0.2% Tricaine, fixed in 4% paraformaldehyde in PBS, cryoprotected in 30% sucrose, and frozen in OCT tissue freezing compound (Fisher). 12 µm-thick sections were cryosectioned for immunostaining. To detect human CNGβ1 or Myc-tagged constructs, tissue sections were permeabilized for 10 min with 0.1% Triton X-100 in PBS, blocked with 5% goat serum and 0.5% Triton X-100 in PBS for 1 hour at 22°C. Specific primary antibodies were diluted 1:2000 in blocking and incubated overnight at 4^°^C. The next day, sections were rinsed in 0.5% PBX and incubated with anti-mouse IgG secondary antibodies conjugated to Alexa Fluor 594 (Jackson ImmunoResearch) and 10 µg/ml Hoechst 33342 (Thermo Fisher Scientific, H3569) in blocking solution for 2 hours at 22°C. Slides were coverslipped using Immu-Mount (Thermo Fisher Scientific).

#### Mouse retinal cross-sections

Posterior eyecups were fixed for 1 hour in 4% paraformaldehyde in PBS, rinsed 3 times in PBS, and embedded in 7.0% low melt agarose (Sigma, A3038). 100 µm cross-sections through the central retina were collected using a vibratome (Leica VT1200S), placed in a 24-well plate, and blocked in 5% goat serum and 0.5% Triton X-100 in PBS for 1 hour at 22°C. Sections were incubated in primary antibody diluted in blocking solution overnight at 4°C, rinsed 3 times in PBS, and incubated with 10 µg/ml Hoechst 33342 and goat or donkey secondary antibodies conjugated with Alexa Fluor 488, 568, or 647 (Jackson ImmunoResearch) in blocking solution for 2 hours at 22°C. Sections were mounted with Immu-Mount and cover-slipped.

At Duke University, images were acquired using a Nikon Eclipse 90i upright microscope equipped with a 100× oil-immersion objective (1.40 NA, Plan Apo VC), A1 confocal scanner controlled by NIS-Elements AR software (Nikon). At the University of Michigan, images were acquired using a Zeiss Observer 7 inverted microscope equipped with a 63× oil-immersion objective (1.40 NA), LSM 800 confocal scanhead controlled by Zen 5.0 software (Zeiss). Manipulation of images was limited to adjusting the brightness level, image size, rotation and cropping using FIJI (ImageJ) and Illustrator (Adobe).

### Protein Deglycosylation Assay

Eyecups from C57BL/6J mice were collected at P21 and homogenized by pestle, followed by sonication in 250 ml of 1% sodium dodecyl sulfate and 1× cOmplete protease inhibitor mixture (Roche, Indianapolis, IN) in PBS. Lysates were cleared at 15,000 rpm for 10 min at 22°C. Total protein concentration was measured using the DC Protein Assay kit (Bio-Rad) and 20 µg of protein was used for each condition. For PNGase treatment, lysate is combined with 2 µL each of Glycoprotein Denaturing Buffer, GlycoBuffer 2 and 10% NP-40, 1µL PNGase F (NEB, P0708L) and H_2_O as necessary to a final volume of 20µL. The reaction is then incubated for 1hr at 37°C. For Endo H (NEB, P0702L) treatment, the same procedure was followed, except that GlycoBuffer 3 was used instead of GlycoBuffer 2. For control experiments, reactions used GlycoBuffer 2 without enzymes. The resulting reactions were terminated by cooling in ice or freezing at −20°C. 5× sample buffer with 100 mM dithiothreitol (DTT) was added to each sample and 5 µg total protein loaded on SDS-PAGE for Western blot.

### Experimental design and statistical analyses

All *in vivo* electroporated mouse retinas were analyzed at postnatal day 21. All transgenic *Xenopus* tadpoles were collected at 14 days post injection (stage 43-45). A minimum of three expressing eyes were analyzed for every DNA construct. Animals of both sexes were used for all experimental models.

## Results

### Domain structure of the rod CNG channel subunits

The domain composition of CNG subunits is shown in Figure 1. Both subunits contain a cytosolic N-terminus, a hexahelical transmembrane core and a cytosolic C-terminus bearing the cGMP-binding domain (Molday and Molday, 1998). The channel’s subunits interact through sites located at the C-terminus of CNGα1 and N-terminus of CNGβ1 (Trudeau and Zagotta, 2002). The N-terminus of CNGβ1 is also responsible for binding calmodulin (Grunwald et al., 1998; Weitz et al., 1998) and peripherin-2 (Ritter et al., 2011; Milstein et al., 2017). The latter is conveyed through a relatively large GARP domain (Sugimoto et al., 1991; Colville and Molday, 1996), which can be sub-divided to a region containing four short proline-enriched repeats responsible for peripherin-2 binding (Ritter et al., 2011; Milstein et al., 2017) and a region of high glutamic acid content. Notably, rods also express two soluble proteins, GARP1 and GARP2, formed through alternative splicing of the *Cngb1* gene and include various fragments of the CNGβ1 N-terminus (Ardell et al., 1995; Ardell et al., 1996; Ardell et al., 2000).

**Figure 1.**
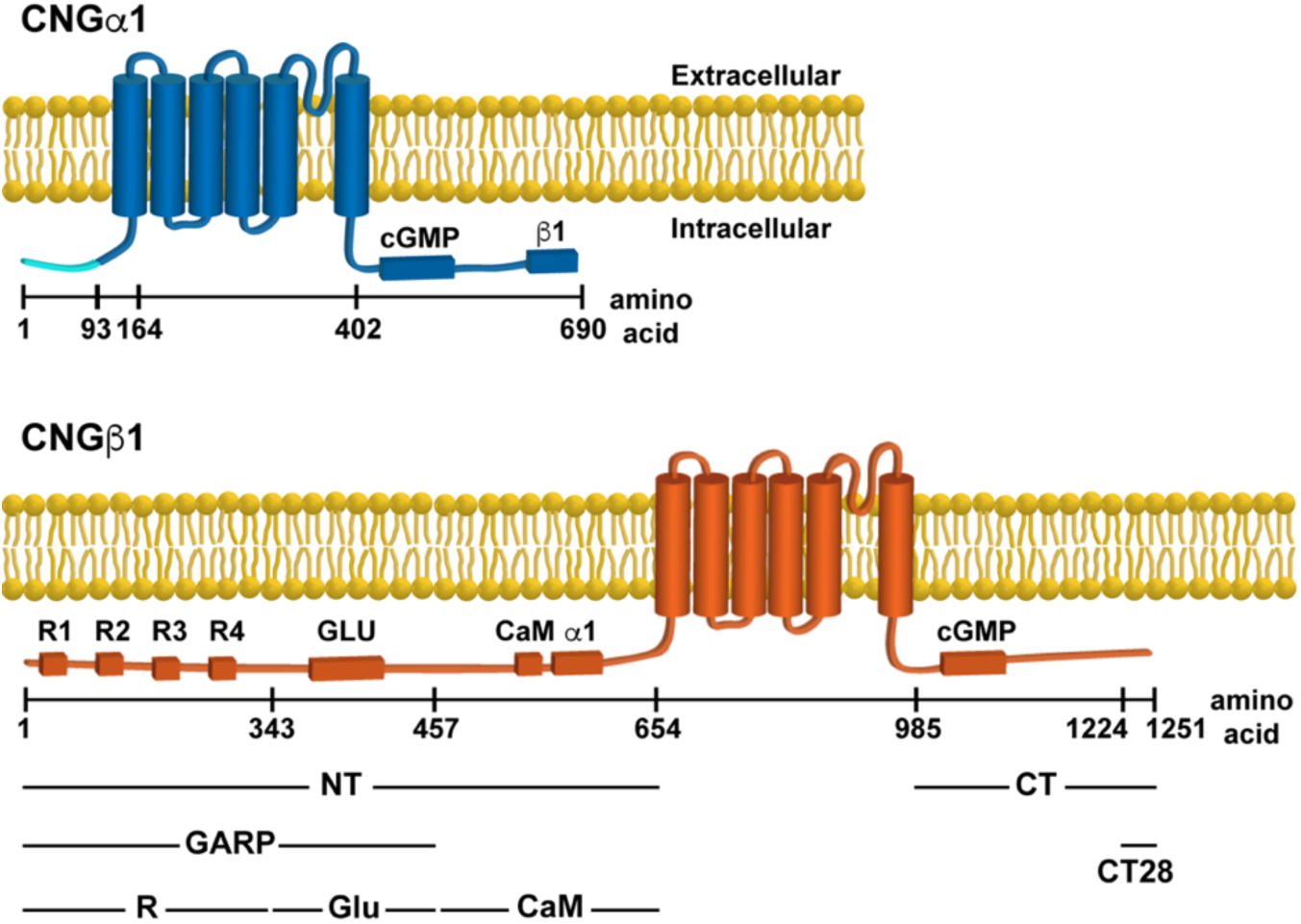
Domain composition of CNGα1 and CNGβ1. Cytosolic domains include: cGMP binding; calmodulin (CaM) binding; CNGα1 (α1) and CNGβ1 (β1) binding; glutamic-acid rich region (GLU) and four proline-rich repeats (R1-R4). The boundaries of CNGβ1-derived constructs expressed in this study shown below the cartoon. The numbering of amino acids corresponds to human protein sequences. Notably, the first 92 amino acids of CNGα1 (marked in light blue) are cleaved during intracellular CNG channel processing (Molday et al., 1991).

### The CNG channel utilizes a conventional trafficking pathway to reach the outer segment

Membrane proteins can utilize two secretory pathways to reach their final destinations: the conventional pathway through both the ER and Golgi and the unconventional pathway directly from the ER. One way to determine which pathway is used by a particular protein is to assess whether its glycosylation is sensitive to Endoglycosidase H (Endo H). N-glycosylation takes place in ER, where high-mannose oligosaccharides are transferred to proteins. This high-mannose N-glycan is sensitive to Endo H cleavage, so proteins exiting from the ER can be fully deglycosylated by this enzyme. If a glycoprotein is transferred and is processed through the Golgi, its oligosaccharide moieties are modified to produce hybrid oligosaccharide chains that are resistant to Endo H cleavage. Accordingly, proteins trafficked through the unconventional secretory pathway are Endo H-sensitive and those using the conventional pathway are Endo H-resistant. In contrast, all N-linked glycoproteins are sensitive to general amidases, like PNGase F.

To understand how outer segment trafficking utilizes these two secretory pathways, we treated mouse retinal lysates with PNGase F and Endo H to examine the glycosylation status of eight outer segment-resident membrane proteins (Figure 2). We found that the electrophoretic mobility of three proteins, CNGβ1, guanylyl cyclase-2 (GC-2) and Rom-1, did not change upon PNGase F or Endo H treatment (Figure 2A), suggesting that they are not N-linked glycoproteins. The other five proteins, CNGα1, rhodopsin, peripherin-2, guanylyl cyclase-1 (GC-1) and ABCA4, had increased mobility upon PNGase F treatment demonstrating that they are N-linked glycoproteins. We also treated retinal lysates with Endo H and found that three proteins, CNGα1, GC-1 and rhodopsin, are resistant to this treatment (Figure 2B), indicating that they are trafficked via the conventional pathway. Whereas this route of outer segment delivery was previously established for rhodopsin (Deretic and Papermaster, 1991), the findings for CNGα1 and GC-1 represent original observations. Notably, the behavior of GC-1 is consistent with our previous result that it is delivered to the outer segment in a complex with rhodopsin (Pearring et al., 2015). Consistent with previous observations (Connell and Molday, 1990; Illing et al., 1997), ABCA4 was completely deglycosylated by Endo H, whereas peripherin-2 displayed a mixed pattern whereby the majority was deglycosylated and a smaller fraction, shown to be associated with Rom-1 (Conley et al., 2019), was not (Figure 2C).

**Figure 2.**
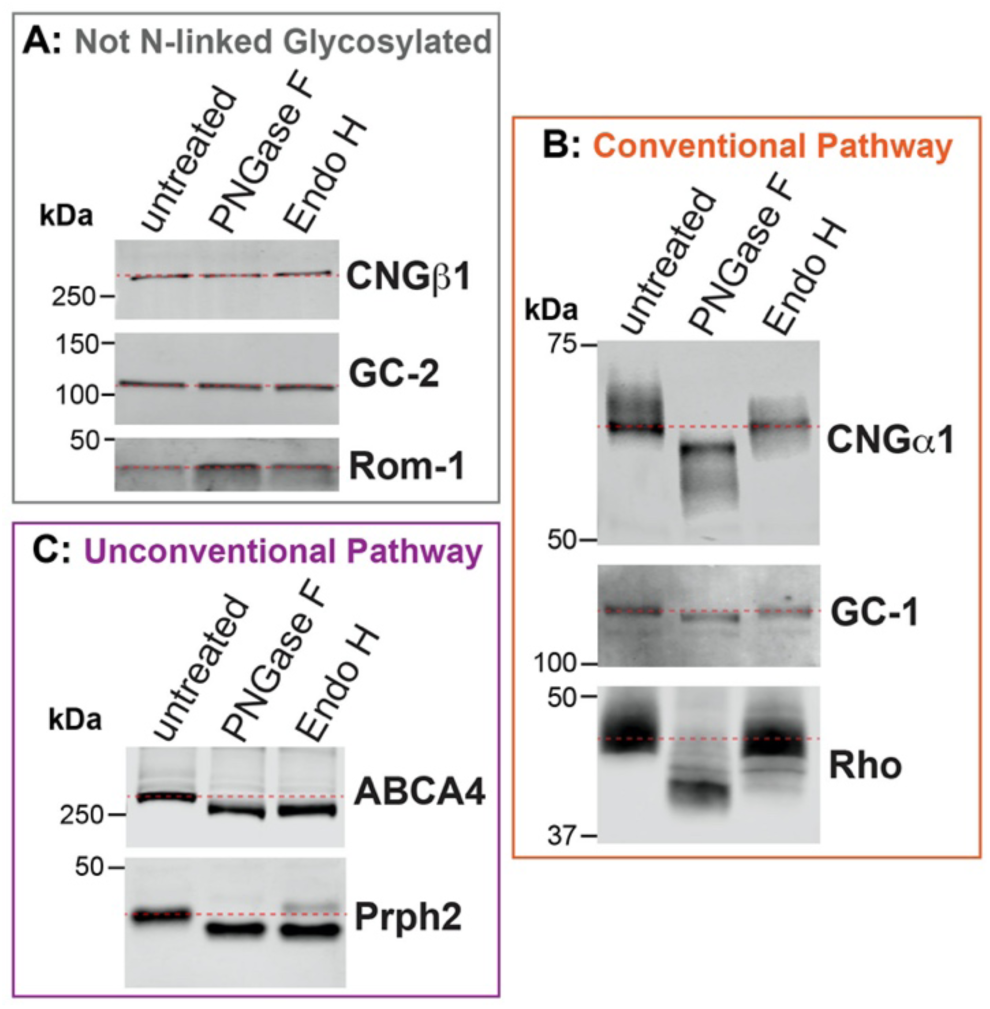
The rod CNG channel is trafficked using the conventional secretory pathway. Mouse retinal lysates (untreated) were incubated with PNGase F or Endo H and analyzed by Western blotting. (**A**) Electrophoretic mobility of non-glycosylated proteins CNGβ1, guanylate cyclase-2 (GC-2) and Rom-1, is unaffected by enzymatic treatments. (**B**) CNGα1, guanylate cyclase-1 (GC-1) and rhodopsin (Rho) are sensitive to PNGase F and resistant to Endo H treatment, indicating their processing through the conventional secretory pathway. (**C**) ABCA4 and peripherin-2 (Prph2) are sensitive to both PNGase F and Endo H treatments, indicating their processing through the unconventional secretory pathway. Dashed red lines are used to mark the positions of untreated protein bands. 10 µg of total protein was loaded in each lane.

These results reveal a peculiar analogy between the trafficking of the peripherin-2/Rom-1 and CNGα1/CNGβ1 complexes. Both complexes utilize the conventional secretory pathway for outer segment delivery and only one subunit in the complex becomes N-link glycosylated en route.

### Outer segment localization of the CNG channel requires subunit pre-assembly and is driven by the CNGβ1-subunit

To investigate the relationship between the CNG channel subunits delivery to the rod outer segment, we first expressed epitope-tagged constructs representing each subunit in both WT and CNGβ1 knockout (*Cngb1-X1*^*−/−*^) mice. The *Cngb1-X1*^*−/−*^ mouse has a targeted deletion of exons 1 and 2 from the *Cngb1* gene resulting in the CNGβ1 knockout (Zhang et al., 2009). It is also characterized by an ∼30-fold reduction in the expression level of CNGα1, suggesting that the cellular stability of CNGα1 relies on its association with CNGβ1. Expression of CNGα1-MYC in WT mice resulted in its predominantly normal outer segment localization (Figure 3A). However, when expressed in *Cngb1-X1*^*−/−*^ mice, CNGα1-MYC was localized in both outer segments and other cellular compartments in a pattern typical for untargeted membrane proteins expressed in mouse rods (Pearring et al., 2014). These observations suggest that CNGβ1 contains targeting information required for the outer segment delivery of the entire channel. Consistently, co-expression of CNGα1-MYC with CNGβ1-FLAG in *Cngb1-X1*^*−/−*^ rods re-directed CNGα1-MYC to the outer segment, along with expressed CNGβ1 (Figure 3B).

**Figure 3.**
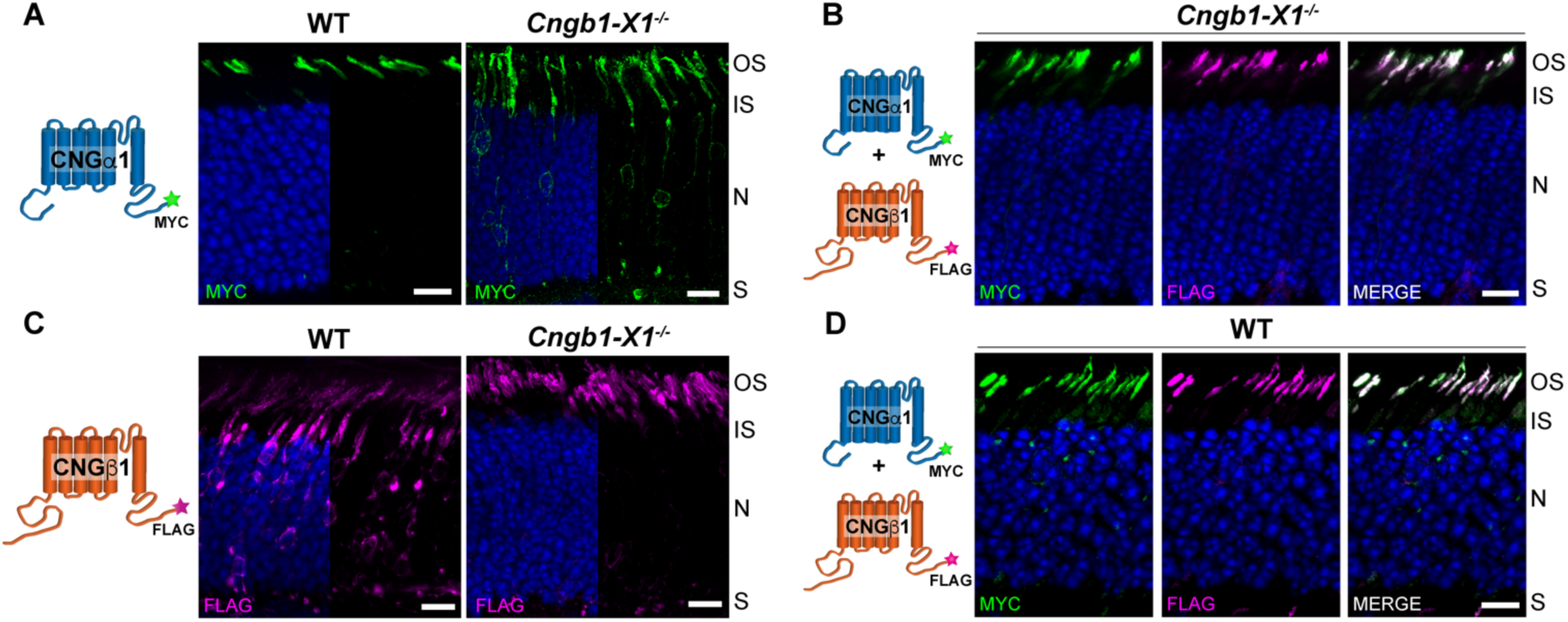
Outer segment localization of the CNG channel requires expression of the CNGβ1 subunit. (**A**) Retinal cross-sections from wild-type (WT) and *Cngb1-X1*^*−/−*^ rods transfected with MYC-tagged, full-length mouse CNGα1. (**B**) *Cngb1-X1*^*−/−*^ rods were co-transfected with CNGα1-MYC and CNGβ1-FLAG. The merged image is shown in the right panel. (**C**) WT or *Cngb1-X1*^*−/−*^ rods transfected with FLAG-tagged, full-length mouse CNGβ1. (**D**) WT rods were co-transfected with CNGα1-MYC and CNGβ1-FLAG. In all panels, retinal sections were stained for each construct using an anti-MYC (green) and/or anti-FLAG (magenta) antibodies as indicated in the panel and nuclei are stained by Hoechst (blue). Cartoon of transfected constructs are depicted on the left. Scale bar, 10 µm. Here and in the following figures, photoreceptor cell layer abbreviations: outer segment (OS), inner segment (IS), nucleus (N), and synapse (S).

There is another important aspect of the data shown in Figure 3A. The fact that all overexpressed CNGα1 is normally targeted in WT rods implies that these cells naturally express more CNGβ1 than is incorporated in the mature channel. This extra CNGβ1 is normally cleared by the cell but can become a part of a mature channel when CNGα1 is overexpressed. Further evidence that the cellular level of the mature CNG channel is determined by the amount of expressed CNGβ1 comes from the behavior of CNGβ1-FLAG expressed in WT and *Cngb1-X1*^*−/−*^ rods. Whereas CNGβ1-FLAG was normally localized to outer segments of *Cngb1-X1*^*−/−*^ rods, a large fraction was retained in the soma of WT rods in a pattern suggesting retention in the biosynthetic membranes (Figure 3C). This suggests that knockout rods contain enough CNGα1 to associate with expressed CNGβ1-FLAG and form a mature channel, but WT rods do not express enough CNGα1 to assemble with both endogenous CNGβ1 and overexpressed CNGβ1-FLAG. Consistent with this interpretation, co-expression of CNGα1-MYC with CNGβ1-FLAG in WT rods resulted in normal localization of both subunits in the outer segment (Figure 3D), showing that increased CNGα1 levels are sufficient to assemble with both endogenous and overexpressed CNGβ1.

### CNGβ1’s C-terminal “RVXP” motif is not required for outer segment localization

Previous studies of the olfactory CNG channel localization in cell culture showed that its ciliary targeting requires a “RVXP” motif on the C-terminus of the CNGβ1b subunit, which is spliced from the same *Cngb1* gene as CNGβ1 expressed in rods (Jenkins et al., 2006; Jenkins et al., 2009). Therefore, we examined whether this sequence in CNGβ1 (RVSP in the mouse) is required for the outer segment CNG channel targeting in rods. We generated a mutant construct in which these four residues were replaced with alanines (CNGβ1_4A_) and expressed it in *Cngb1-X1*^*−/−*^ rods. The outer segment localization of this mutant was indistinguishable from the WT CNGβ1 control (Figure 4A), demonstrating that the “RVXP” motif does not confer outer segment targeting. A similar result was obtained upon transgenic expression of the corresponding mutant of human CNGβ1 subunit in *Xenopus* rods. Both outer segment targeting and subcellular localization of the mutant construct to the outer segment plasma membrane were normal (Figure 4B). Notably and unlike in WT mice, we did not observe CNGβ1 retention in the biosynthetic membranes upon overexpression, which may be explained by differences in relative overexpression levels and/or cross-species differences in the clearance efficiency of unprocessed proteins.

**Figure 4.**
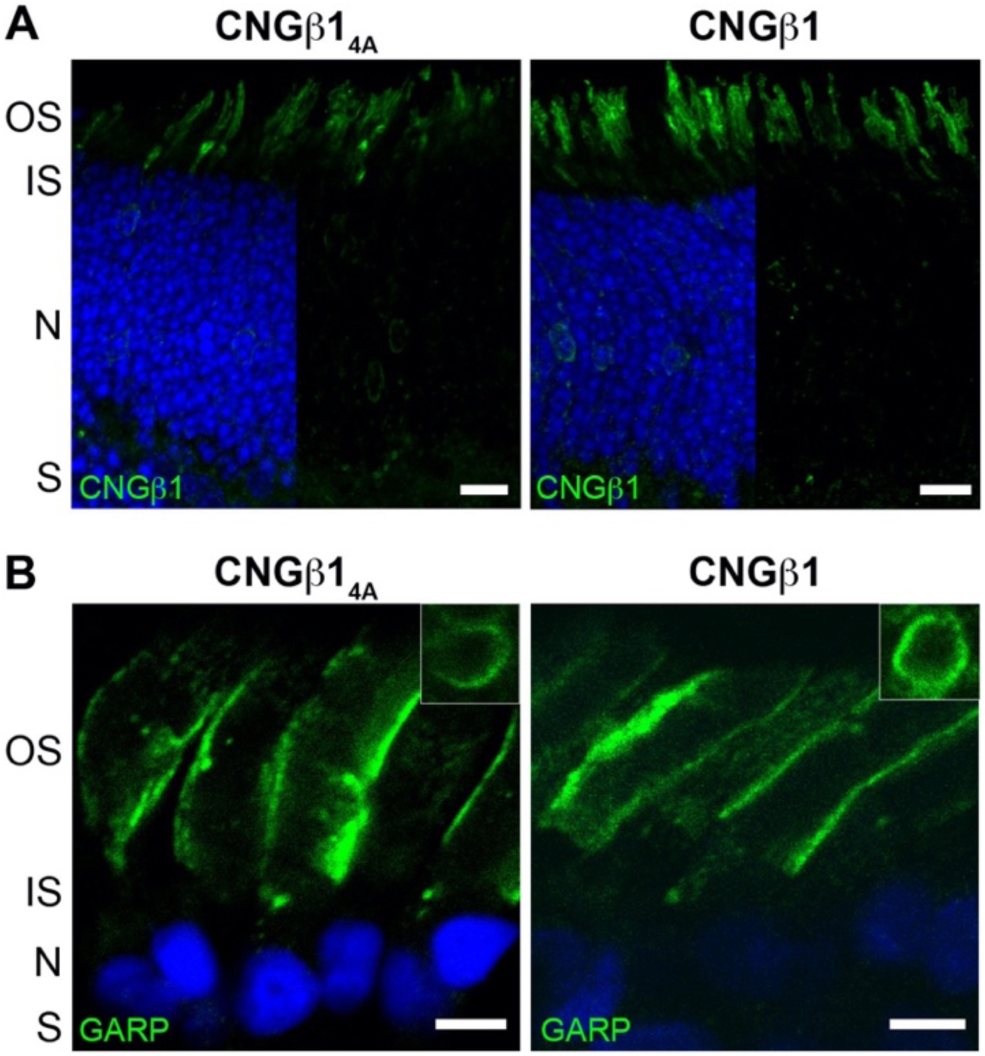
The C-terminus “RVSP” motif is not involved in CNG channel localization to the plasma membrane of the outer segment. (**A**) *Cngb1-X1*^*−/−*^ rods were transfected with untagged mouse CNGβ1_4A_ mutant or CNGβ1 control constructs. Retinal cross-sections were stained for each construct using an anti-CNGβ1 (green) antibody. Scale bar, 10 µm. (**B**) Transgenic *Xenopus* rods expressing full-length human CNGβ1_4A_ mutant or CNGβ1 control. Retinal cross-sections were stained for each construct using an anti-GARP (green) antibody. Scale bar, 5 µm. Nuclei are counterstained with Hoechst (blue).

### Outer segment targeting of CNGβ1 is mediated by its N-terminus

In the next set of experiments, we undertook a search for the outer segment targeting and sorting signal(s) within CNGβ1. We first expressed the cytosolic C-terminus of human CNGβ1 in transgenic *Xenopus* (CT construct in Figure 1). To ensure membrane-association while preserving its relative membrane topology, this construct was fused with an N-terminal single-pass transmembrane domain, subcloned from the activin receptor. However, the resulting TM-CNGβ1_CT_ construct was retained in the biosynthetic membranes (Figure 5A) precluding us from analyzing its targeting capacity. To circumvent this problem, we appended the CT sequence to a construct containing YFP and the untargeted lipidation sequence of rhodopsin (YFP-Rho_CTΔ5_), a strategy previously applied to achieve membrane attachment of C-terminal protein fragments (Tam et al., 2004; Salinas et al., 2013) (YFP-RhoCTΔ5-CNGβ1_CT_; Figure 5B). However, expression of the YFP-RhoCTΔ5-CNGβ1_CT_ construct resulted in a pattern indicative of a soluble protein, suggesting that the long full-length CT construct prevented posttranslational lipidation of the rhodopsin C-terminus. Therefore, we resorted to testing a short C-terminal sequence of 28 amino acids previously suggested to contain outer segment targeting information (Kizhatil et al., 2009). Somewhat unexpectedly, when we expressed the YFP-Rho_CTΔ5_-CNGβ1_CT28_ its subcellular distribution was essentially identical to that of the base YFP-Rho_CTΔ5_ construct (Figures 5C,D). Both constructs followed a “default” distribution pattern for untargeted membrane-associated proteins, in which the majority is localized in the outer segment but appreciable fractions are also present in the plasma membrane of the cell body and the synaptic region (Baker et al., 2008).

**Figure 5.**
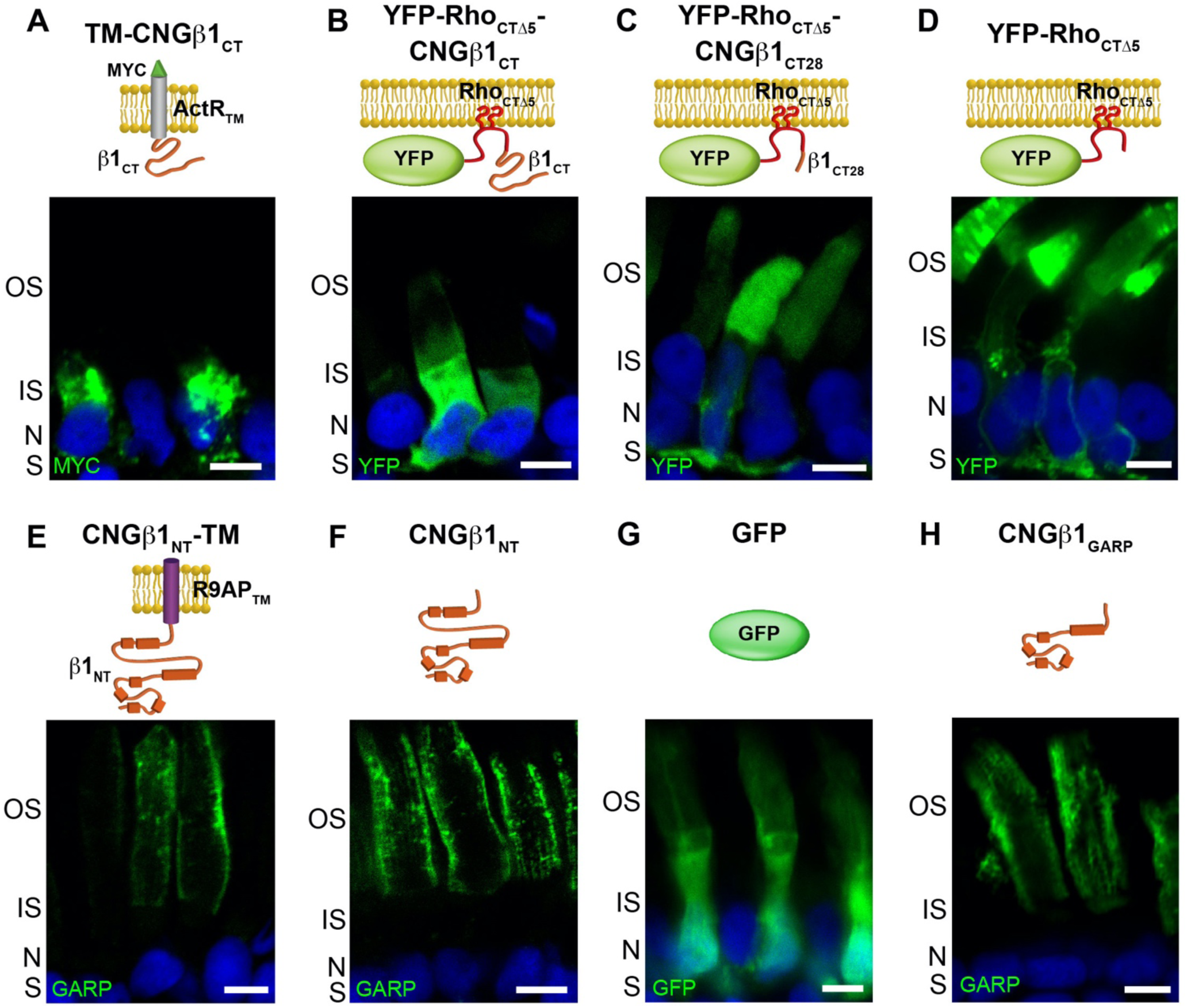
CNGβ1 N-terminal GARP domain is confined to the rod outer segment. Retinal cross-sections from transgenic *Xenopus* rods expressing (**A**) membrane-anchored TM-CNGβ1_CT_, (**B**) lipidated YFP-Rho_CTΔ5_-CNGβ1_CT_, (**C**) lipidated YFP-Rho_CTΔ5_-CNGβ1_CT28_, (**D**) lipidated YFP-Rho_CTΔ5_, (**E**) membrane-anchored CNGβ1_NT_-TM, (**F**) soluble CNGβ1_NT_, (**G**) soluble GFP and (**H**) soluble CNGβ1_GARP_ constructs. Cartoon of transfected constructs are shown above their corresponding panel. Constructs that include N-terminal of human CNGβ1 were detected using an anti-GARP antibody. Nuclei are counterstained with Hoechst (blue) and scale bar is 5 µm in all panels.

Next, we focused on the targeting properties of the cytosolic CNGβ1 N-terminus (NT construct in Figure 1). Its proper membrane attachment and topology were established by fusing with the single-pass transmembrane domain from R9AP lacking specific targeting information (Baker et al., 2008; Pearring et al., 2014). The resulting CNGβ1_NT_-TM construct was localized exclusively to the outer segment, suggesting that it bears sufficient targeting information (Figure 5E). However, when we expressed the same construct as a soluble protein (CNGβ1_NT_), it was also localized to the outer segments with the same staining pattern (Figure 5F). The latter is inconsistent with the well-described distribution of soluble proteins, such as GFP, located predominantly in the cytosol-rich inner segment ((Tam et al., 2000; Najafi et al., 2012); Figure 5G). The distribution of CNGβ1_NT_ resembles of soluble GARP1/2 proteins comprised predominantly from the parts of the CNGβ1 N-terminus (Hüttl et al., 2005; DeRamus et al., 2017) and suggests that its outer segment localization is conveyed through an interaction with another outer segment-resident protein, such as peripherin-2 located at the disc rims or CNGα1 located in the outer segment plasma membrane (Poetsch et al., 2001; Ritter et al., 2011). We then expressed a soluble construct encoding CNGβ1’s GARP domain (CNGβ1_GARP_; Figure 5H) and found that it is distributed like the entire N-terminus. Because the interaction site with CNGα1 resides outside the GARP domain (Figure 1), this result favors peripherin-2 as the interacting partner for these constructs. This was further supported by the analysis of tangential sections cut across rod outer segments expressing soluble N-terminal constructs (Figure 6). Both CNGβ1_NT_ and CNGβ1_GARP_ displayed a flower petal-like pattern, which corresponds to invaginating incisures at the disc rims. This pattern is similar to that of peripherin-2 but is different from that of CNGβ1, which lacks immunostaining of the disc incisures (Figure 6).

**Figure 6.**
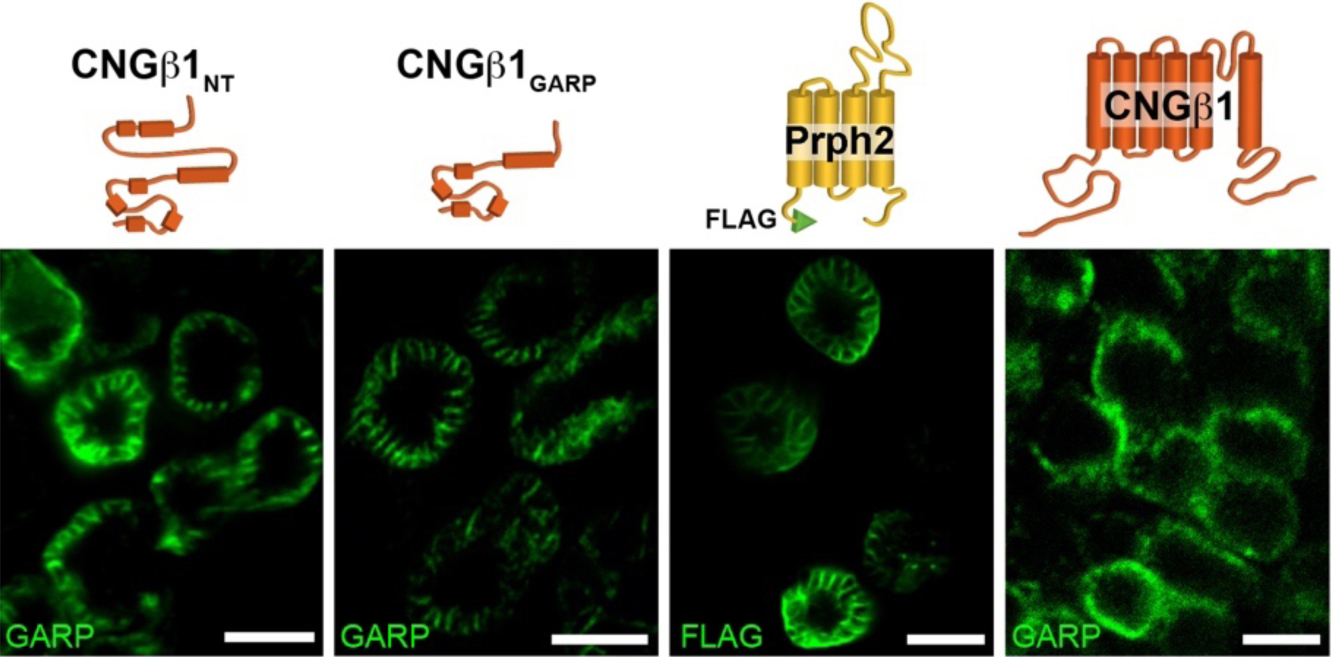
CNGβ1’s N-terminal GARP domain localizes to disc incisures within rod outer segments. Tangential sections through *Xenopus* rod outer segments expressing soluble CNGβ1_NT_ and CNGβ1_GARP_; FLAG-tagged full-length bovine peripherin-2 or untagged full length CNGβ1. Cartoon of transfected constructs are shown above their corresponding panel. Constructs that include N-terminal of human CNGβ1 were detected using an anti-GARP antibody. Scale bar, 5 µm in all panels.

### CNGβ1’s GARP domain encodes both outer segment targeting and peripherin-2 interaction site

Demonstrating that the GARP domain of CNGβ1 binds to the disc rims might suggest that the CNG channel uses an interaction with peripherin-2 for outer segment delivery rather than utilizing its own targeting signal. To explore this possibility, we expressed smaller fragments of the CNGβ1 N-terminus – CNGβ1_CaM_, CNGβ1_R_ and CNGβ1_Glu_ (Figure 1) – as both soluble and membrane-anchored proteins. Soluble CNGβ1_CaM_ was localized predominantly in the inner segment, suggesting that it is not engaged in any outer segment interactions, whereas the corresponding membrane-anchored construct (CNGβ1_CaM_-TM) was retained in the biosynthetic membranes (Figure 7A). Soluble CNGβ1_R_ was biased to the outer segment where it tended to accumulate at the periphery (Figure 7B). However, a portion of this construct was also found in the cell body and synaptic termini, suggesting that its affinity for periphern-2 is not as high as that of longer N-terminal constructs (Figure 5E,G). Membrane-anchored CNGβ1_R_-TM was successfully exiting the biosynthetic membranes and delivered to both inner and outer segments (Figure 7B), indicating that it lacks specific outer segment targeting information. Within the outer segment, CNGβ1_R_-TM was located at disc incisures similar to longer N-terminal constructs, as evidenced from the flower petal-like pattern seen in tangential sections (Figure 7B). Together, these data suggest that the CNGβ1_R_ region is sufficient for peripherin-2 interaction, but not for specific outer segment targeting.

**Figure 7.**
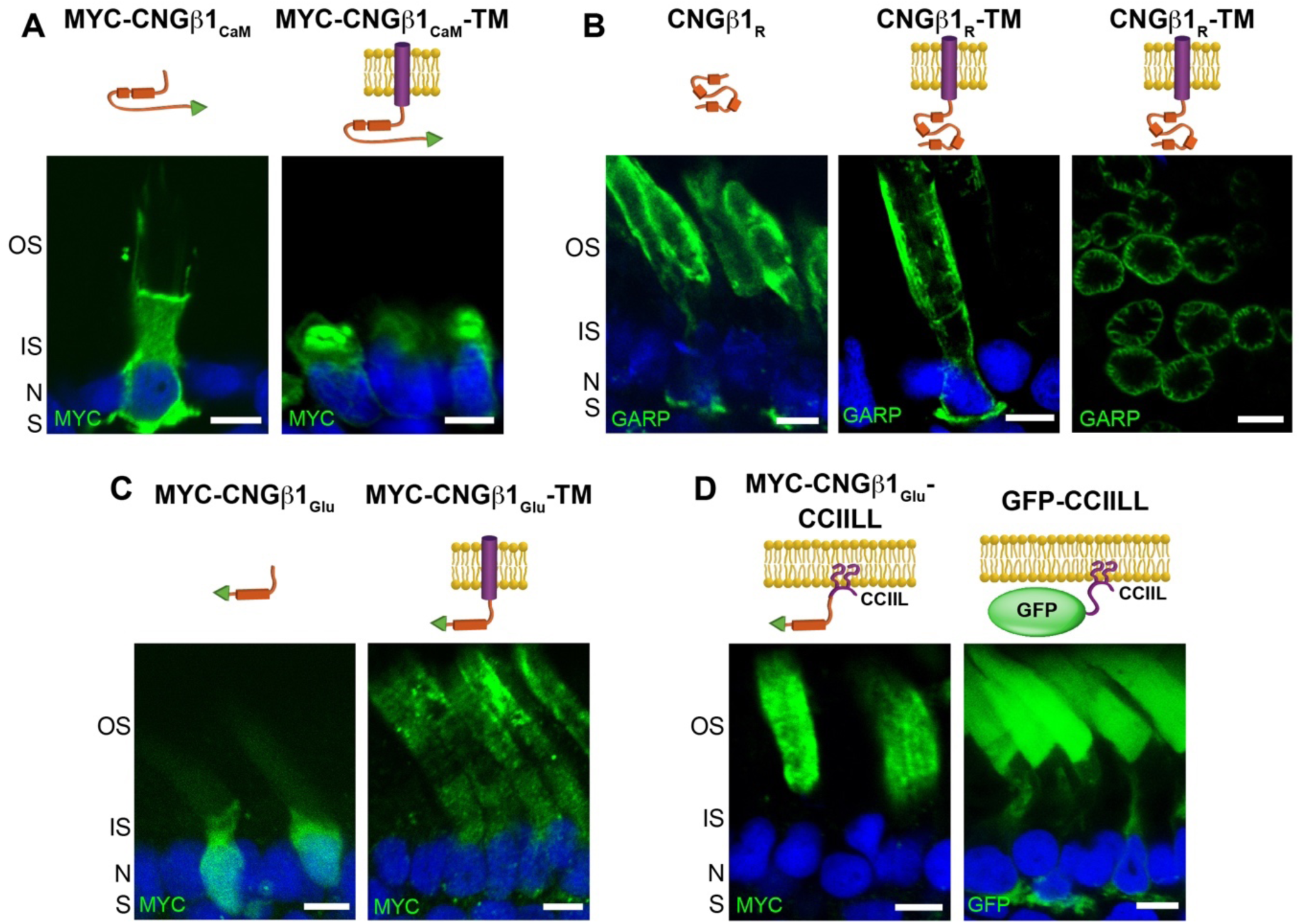
CNGβ1’s GARP domain contains separate sites for peripherin-2 binding and outer segment targeting. Retinal cross-sections from transgenic *Xenopus* rods comparing localization of (**A**) soluble and membrane-anchored CNGβ1_CaM_; (**B**) soluble and membrane-anchored CNGβ1_R_ and a tangential image of outer segment’s expressing CNGβ1_R_-TM; (**C**) soluble and membrane-anchored CNGβ1_Glu_; and (**D**) CNGβ1_Glu_ or GFP fused to a C-terminal CCIIL double lipidation anchor (**D**). Cartoon of transgenic constructs are shown above their corresponding panel. Constructs that include the N-terminal of human CNGβ1 were detected using an anti-GARP antibody. All other constructs were N-terminally tagged with the MYC epitope and detected using an anti-MYC antibody. Scale bar, 5 µm. Nuclei are counterstained with Hoechst (blue).

We next analyzed CNGβ1_Glu_ constructs (Figure 7C). Soluble CNGβ1_Glu_ behaved as a typical cytosolic protein with diffuse staining within the photoreceptor cytoplasm, suggesting the lack of specific outer segment interactions. The membrane-anchored CNGβ1_Glu_-TM was found in both inner and outer segments. However, its inner segment fraction was located inside the cell and not within the surrounding plasma membrane. This suggests that only a portion of this construct is properly processed by the biosynthetic membranes, precluding us from determining whether CNGβ1_Glu_ may contain an outer segment targeting signal. To avoid protein misprocessing, we expressed CNGβ1_Glu_ as a lipidated construct by adding a C-terminal CCIIL sequence for double lipidation (palmitoylation and geranylgeranylation) previously introduced in similar studies (Tam et al., 2004; Baker et al., 2008; Pearring et al., 2014) (Figure 7D). We have found that CNGβ1_Glu_-CCIIL was localized exclusively in the outer segments, whereas the control untargeted GFP-CCIIL construct displayed a typical default pattern with an appreciable amount localized in other parts of the rod cell. This result shows that CNGβ1_Glu_ contains a specific outer segment targeting signal. Within the outer segment, CNGβ1_Glu_-CCIIL was evenly distributed across all membranes without showing any bias toward periphery. Combined with the behavior of the soluble CNGβ1_Glu_ construct, this suggests that this region of CNGβ1 is not sufficient for binding to peripherin-2.

## Discussion

Protein targeting to the light-sensitive photoreceptor outer segment is still far from being understood and remains a subject of active investigation. Except for a few proteins shown to be delivered within larger complexes, such as GC-1 and PRCD associated with rhodopsin (Pearring et al., 2015; Spencer et al., 2016) or RGS9 and Gβ5 associated with R9AP (Hu and Wensel, 2002; Keresztes et al., 2004; Krispel et al., 2006; Gospe et al., 2011), most membrane proteins have targeting information encoded in their sequences. Amongst them, the CNG channel is particularly interesting because it needs to be both delivered to the outer segment and then sequestered to the plasma membrane surrounding the discs. In this study, we demonstrated that the rod outer segment delivery of this channel requires pre-assembly of its constituent subunits upon their biosynthesis and that the targeting signal is encoded within the glutamic acid-rich region of CNGβ1’s GARP domain. Whereas acidic clusters have previously been associated with packaging membrane proteins into clathrin-coated vesicles (Navarro Negredo et al., 2017), to our knowledge, this is the first time an acidic cluster has been implicated in ciliary targeting of a membrane protein. This targeting signal is sufficient for outer segment localization but not subsequent outer segment plasma membrane sequestration, suggesting that the final sorting of the CNG channel to the outer segment plasma membrane occurs independently of its initial targeting.

We further showed that the cellular level of the mature CNG channel in rods is determined by the expression level of CNGβ1. This follows the pattern in which the total amount of a mature multi-subunit protein complex is determined by the expression level of one of its subunits. The best studied example in photoreceptors is the RGS9-Gβ5-R9AP GTPase activating complex for transducin, whose cellular content is set by the expression of R9AP, while the fractions of other subunits expressed in excess of R9AP are efficiently cleared by the cell (Keresztes et al., 2004; Krispel et al., 2006).

Another aspect of outer segment trafficking explored in this study is that some proteins, such as rhodopsin, use the conventional secretory pathway through the ER and Golgi (Deretic and Papermaster, 1991; Deretic et al., 1998; Murray et al., 2009), while others, such as ABCA4, use the unconventional pathway bypassing the Golgi (Illing et al., 1997; Tsybovsky et al., 2011). Peripherin-2, known to interact with the CNG channel in outer segments, is also delivered through the unconventional pathway (Connell and Molday, 1990). The exception is a fraction of peripherin-2 engaged in the hetero-tetrameric complex with Rom-1, which is diverted into the conventional pathway (Zulliger et al., 2015; Conley et al., 2019). The latter is species-specific: it was documented for mice, yet the entire pool of peripherin-2 in bovine and *Xenopus* is delivered via the unconventional pathway (Connell and Molday, 1990; Tian et al., 2014).

Our new data show that the CNG channel is delivered to the outer segment via the conventional pathway, which may suggest that the CNG channel and peripherin-2 are trafficked to this compartment independently of one another, particularly because the entire pool of peripherin-2 was shown to utilize the unconventional pathway in *Xenopus* (Tian et al., 2014). However, another *Xenopus* study employing fluorescence complementation assays concluded that the interaction between the CNG channel and peripherin-2 first takes place in the inner segment (Ritter et al., 2011). Furthermore, CNGβ1 is completely absent from the outer segment membrane material of peripherin-2 knockout mice despite all other known outer segment proteins, including CNGα1 and Rom-1, being found in this preparation (Spencer et al., 2019). This argues that the CNGβ1 cannot be delivered to the outer segment without peripherin-2 and they ought to interact before reaching this destination. A possible trafficking mechanism reconciling all of these observations was proposed in a recent study suggesting that, at least in cones, peripherin-2 is trafficked through the late endosome before entering the outer segment (Otsu et al., 2019). Likewise, the CNG channel might pass through endosomes, based on its mislocalization from the outer segment in conditional double knockout mice lacking the endocytic adaptor proteins Numb and Numb-like in rods (Ramamurthy et al., 2014). The authors further showed that, in cell culture, Numb binds and re-directs CNGα1 to endosomes. Therefore, it is conceivable that the CNG channel and peripherin-2 are first trafficked to the inner segment endosomes using the conventional and unconventional secretory pathways, respectively, and then are delivered to the outer segment as a complex.

If the CNG channel and peripherin-2 are in fact delivered to the outer segment as a complex, then how do they next segregate into two different membrane sub-compartments – the discs and the plasma membrane? It logical to assume that they segregate upon the remodeling of outer segment membranes during the process of disc enclosure (Spencer et al., 2020a), after which they remain connected in-trans forming a link between the disc rim and the plasma membrane. However, no direct evidence for this mechanism have been obtained so far. It also remains unknown whether any additional proteins, such as the Na/Ca-K exchanger associated with the CNG channel, are involved in the channel’s plasma membrane segregation. It has been proposed that this process involves a cytoskeletal protein, ankyrin-G (Kizhatil et al., 2009). However, subsequent proteomic studies have not identified any ankyrin-G peptides in the outer segment (Skiba et al., 2013; Spencer et al., 2019) challenging the existence of ankyrin-G in this subcellular compartment. Thus, elucidating the exact outer segment trafficking route of the CNG channel and the mechanism responsible for its subsequent sorting into the outer segment plasma membrane remain the challenges for future studies.

## Acknowledgements

This work was supported by the NIH grants EY025732 (JNP), EY012859 (VYA), EY030451 (VYA), EY007003 (University of Michigan), EY005722 (Duke University), Matilda E. Ziegler Research Award (JNP) and Research to Prevent Blindness.

## Notes

Conflict of Interest Statement: The authors declare no competing financial interests.

### Competing Interest Statement

The authors have declared no competing interest.

